# Neurofeedback enhances a neural signature of selective attention to speech in cocktail-party settings

**DOI:** 10.64898/2026.05.29.728767

**Authors:** Ram K. Pari, Marina Inyutina, Hoang Lam Thanh Hoang, Camille Dedies, Manuela Jaeger, Stefanie Enriquez-Geppert, Mathieu Marx, Stefan Debener, Christoph S. Herrmann, Benedikt Zoefel

**Affiliations:** Université de Toulouse, CNRS, Centre de Recherche Cerveau et Cognition (CerCo), Toulouse, France; CHU Toulouse Purpan, Toulouse, France; Neuropsychology Lab, Department of Psychology, University of Oldenburg, Oldenburg, Germany; Department of Clinical and Developmental Neuropsychology, University of Groningen, The Netherlands; Cluster of Excellence “Hearing4All”, Carl von Ossietzky University, Oldenburg, Germany; Research Center Neurosensory Science, Carl von Ossietzky University, Oldenburg, Germany; Experimental Psychology Lab, Department of Psychology, Carl von Ossietzky University, Oldenburg, Germany

## Abstract

Understanding speech in noisy situations is challenging and often fails due to attentional rather than sensory deficits. We report here that neurofeedback can enhance a neural signature of selective attention to speech in a cocktail-party setting. Neural responses to speech were quantified in participants’ electroencephalogram (EEG) while they attended to one of two audiobooks, presented simultaneously. After each 22s-segment of speech, the participants’ N1 component was extracted from the temporal response function (TRF). The N1 component is known to be sensitive to attention. During a ∼49-min training session, N1 amplitude values were displayed visually to participants so that they could learn to strengthen their neural responses to target speech and minimise their responses to distracting speech. During neurofeedback training, we found an enhanced N1 component in the response to target audiobooks that was specific to EEG channels used to provide feedback, and not present in a control group that received sham feedback. At right-lateralised fronto-central channels, enhanced N1 components correlated with improvements in a measure of speech comprehension (multiple-choice content questions). These results indicate that neural responses to speech can be regulated through neurofeedback and open up new possibilities to train attentional listening in populations struggling to understand speech in noise.

## Introduction

Speech perception is a crucial aspect of everyday life, but becomes challenging in noisy scenarios such as in a busy restaurant (Cherry, 1953; King & Walker, 2020). This challenge becomes more prominent during ageing, when comprehension of speech in noise declines rapidly (Lin et al., 2013). This decline is not necessarily caused by actual hearing loss (Füllgrabe et al., 2015) but can reflect a reduced ability to suppress distracting speakers or noise, suggesting a contribution of attentional deficits (Gillingham et al., 2018; Tun et al., 2009). At the same time, suppression of background noise is a task that current hearing aids often fail at (Dong et al., 2024; Wu et al., 2019). It is therefore important to find alternative ways of improving attention to speech in noise. We report here a successful modulation of a neural signature of attention to speech in noise through neurofeedback.

This approach required a reliable measure of attention to speech in noise, which recent electroencephalography (EEG) research has made possible. Temporal Response Functions (TRFs) allow the modelling of the neural response to a continuous stimulus like speech (Crosse et al., 2016), unlike Event-Related Potentials (ERPs) that are typically computed for individual or discrete events. Recent work found that some TRF components estimated for responses to continuous speech are modulated by selective attention, in particular the “N1” (∼80-150 ms) (Jaeger et al., 2025; Kurthen et al., 2021; Orf et al., 2023; Vanthornhout et al., 2019), resembling attentional effects established for ERPs (Herrmann & Knight, 2001; Rusiniak et al., 2013).

In the current study, participants were provided with TRF-based neurofeedback to train their attention to speech in noise. Neurofeedback (NF) is a method of training to modulate one’s own neural activity. Traditionally, this form of training involves presenting the neural activity of interest to the individual through visual or acoustic stimulation, so participants can use the feedback to learn to regulate their neural activity (Enriquez-Geppert et al., 2017). Several studies have demonstrated the efficacy of neurofeedback to modulate spectral power and ERP components (Arvaneh et al., 2019; Hayashi et al., 2022; Ruddy et al., 2018; Zoefel et al., 2011).

Neurofeedback is very rarely used in combination with TRFs and attention to speech. Kim et al. (2021) employed a 4-session neurofeedback training protocol that was based on EEG responses to words embedded in a non-speech background. Compared to a placebo group, they reported enhanced performance in a speech-in-noise task, as well as stronger evoked responses to target words. Although that result suggests that attention to speech in noise can be trained with neurofeedback, that study only used two isolated words (“up” and “down”). Combined with a small sample size (N=10), it therefore remained unclear whether similar effects can be achieved during continuous speech with competing speakers.

Two recent studies provided participants with neurofeedback on how well attended or ignored speech can be extracted (“decoded”) from their EEG. Jaeger et al. (2020) used cross-correlation for attentional decoding and presented decoding accuracy to participants. Neurofeedback was given exploratorily to test the feasibility of attentional decoding in a complex listening situation, but neurofeedback benefit was not studied. Haro et al. (2025) combined TRFs with neurofeedback. They displayed decoding accuracy for both target and ignored speech streams to participants as feedback. They found reduced neural responses to ignored speech in the second half of neurofeedback training, but no effects on responses to target speech. Moreover, reduced processing of ignored speech was not accompanied by changes in speech comprehension, and how neurofeedback training affects the underlying TRFs was not reported.

Our neurofeedback approach differs from those described in that it relies on a specific component of the TRF that is modulated by attention to speech – the N1. We assumed that this approach links the feedback more closely to attentional processes and is therefore suitable for attentional training. We tested this idea in normal-hearing young adults that completed one session (∼49-min) of neurofeedback. We used natural speech to improve ecological validity. We hypothesised that N1 neurofeedback training allows participants to improve their attentional focus, leading to changes in N1 amplitude relative to baseline sessions without training, and to a Sham Feedback (SF) group. We also expected corresponding improvements in a measure of attentive listening to speech, consisting of multiple-choice content questions, which was acquired in parallel.

## Methods

### Study Participant details

Given the novelty of the design, it was difficult to predict effect sizes with precision. We therefore collected data from as many participants as it was possible in practice. This sample size (total N=56; 38 females) exceeds that reported in typical neurofeedback studies (Treves et al., 2025). The participants were aged between 18 and 35 years (mean age 24 years). One participant was excluded due to issues with data recording. All participants had a native-level proficiency in French and reported normal hearing (in the absence of noise) and no history of neurological disorders. All participants were informed about methods applied and gave written consent to participate in the study. They were also offered compensation of 35 Euros for their study participation. The study was approved by the ethics board, Comité de Protection des Personnes (CPP, protocols 2020-A02707-32 and 2024-A01919-38). Participants were randomly assigned to either a neurofeedback (NF, n=28 participants) or sham feedback (SF, n=27 participants) group based on a predetermined randomisation table, ensuring that the study was double-blinded. Participants were not aware of the possibility of receiving sham feedback; however, they were debriefed after the experiment.

### Stimulus material

Participants were presented with two concurrently played audiobooks in French: one spoken by a male speaker (“The Belton Estate” by Anthony Trollope) and another by a female speaker (“Sense and Sensibility” by Jane Austen). All processing of the audiobooks was done using Audacity (2024). Both audiobooks were cut into snippets of 22s, with a 1s fade-in and fade-out interval. The audio snippets were sampled at 44100 Hz with a 16-bit resolution. The sound level across both audiobooks was matched to a perceptual loudness of -23dB LUFS. Any silent periods were trimmed down to a maximum duration of 500ms.

### Study design

The completed checklist for the reporting and experimental design of clinical and cognitive-behavioural neurofeedback studies (CRED-nf) can be found in the supplementary materials (Ros et al., 2020).

**Fig. 1A** gives an overview of the study design that consisted of several experimental steps, described below in detail. Participants were seated in a dimly lit room, separated from the experimenter by means of a sound-dampening wall. All experimental stimuli were presented using MATLAB 2019a (The MathWorks Inc., 2019) and Psychtoolbox (Brainard, 1997). All auditory stimuli were played through a pair of JBL Control 1Pro speakers, one placed in front of the participant (0°) and another placed behind the participant (180°), driven by an RME Fireface 802 FS sound card. Target speech (that participants were asked to attend to) was always presented from the 0° speaker, and distracting speech (if applicable) from the 180° speaker. The same soundcard was used to send triggers to the EEG system to ensure synchronisation between the presented auditory stimuli and the EEG.

**Figure 1.**
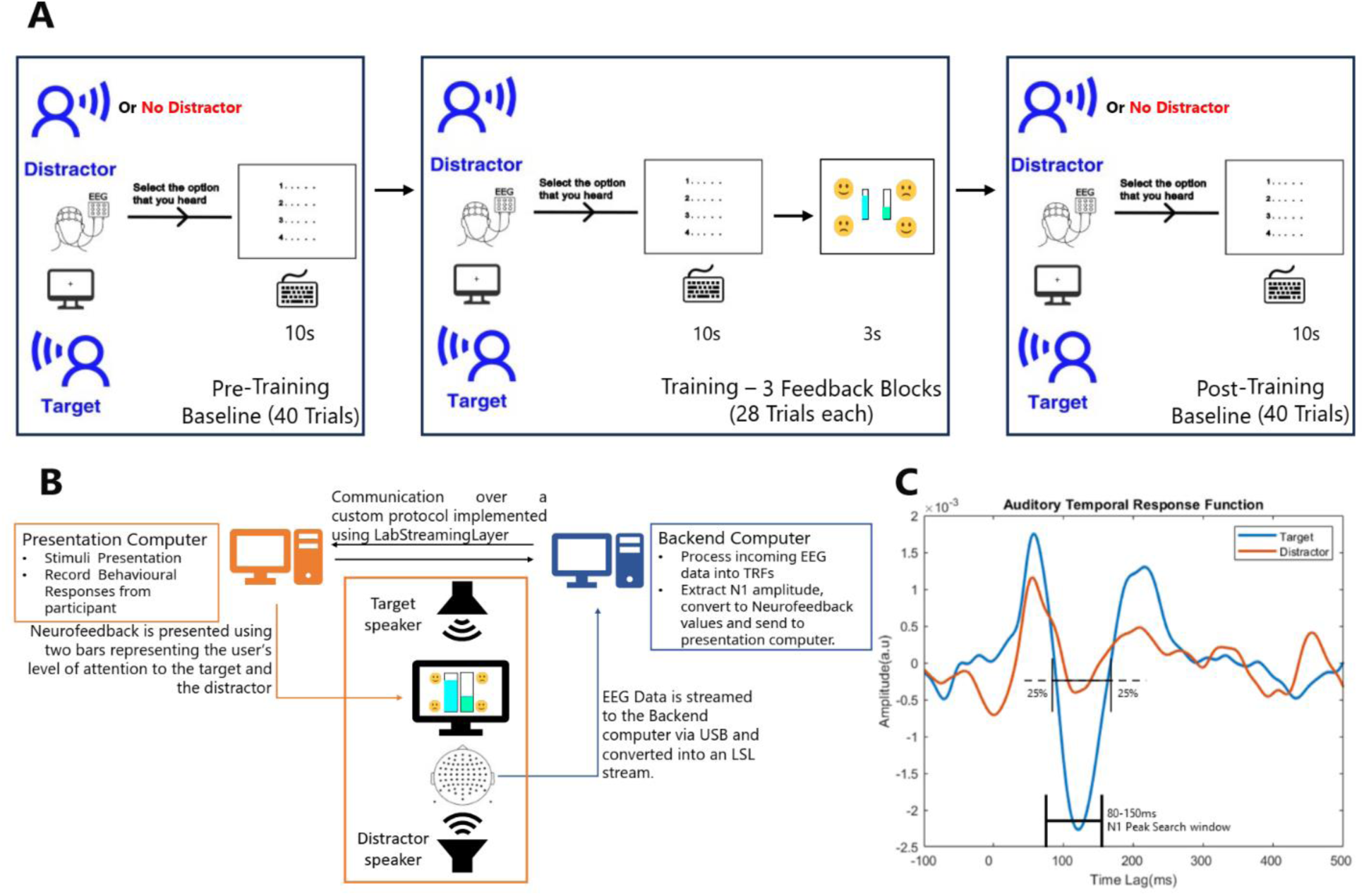
Experimental setup and Algorithm for N1 extraction from Temporal Response Functions (TRFs). **A**. Neurofeedback training was divided into Baseline blocks (pre- or post-training measures) and Feedback blocks (with neurofeedback or sham feedback, depending on the group). During all blocks, participants were instructed to pay attention to a target audiobook, presented at 0°). After each 22-s segment of speech, participants selected one out of four four-word phrases that they believed was part of the preceding segment. A distractor audiobook was presented at 180° in all Feedback trials, and in half of the Baseline trials. Only in Feedback blocks, feedback was presented to the participant in the form of two visual bars, one for the Target and the other for the Distractor. Participants were instructed to increase the displayed value for the Target and reduce it for the Distractor, and instructed that they can achieve this goal by paying attention to the target audiobook, and by ignoring the other audiobook, respectively. Prior to neurofeedback training, participants completed a short volume calibration (not shown). **B**. During each 22-s audio segment, EEG data was collected and processed by a Backend computer to extract TRFs estimated from neural responses to target and distracting audiobooks. The N1 component was then extracted separately for each audiobook and converted to feedback (see Online Data Processing) through a Presentation Computer. **C**. To estimate individual time ranges for the TRF’s N1 component, the largest negative peak was identified first within a search window (80-150ms) and the nearest positive peaks (>10ms) preceding and following the negative peak, respectively. The individual N1 time range was defined as ±25% of the distance between the negative peak and the positive peak on each side, yielding a window centred on the negative peak. N1 time ranges were identified separately for each audiobook (male/female) speaker and condition (Target / Distractor). For each 22-s trial, feedback was calculated as the average N1 in the individual time range, expressed as a percentile relative to the previously obtained values (see Online Data Processing). For illustration, the TRFs shown were obtained from four pilot participants that did not receive neurofeedback and were not included in the final data analysis.

### Adaptive staircase volume calibration

During all experimental steps, audiobooks were presented at +60 dB relative to the estimated individual hearing threshold. This ensured both a comfortable presentation level and approximately similar loudness perception across participants. For this purpose, participants were first asked to perform an adaptive staircase prior to the main experiment. The Palamedes Toolbox was used for this procedure (Prins & Kingdom, 2018). A 1-second-long speech-shaped noise was created with a spectrum equal to the average of all the audio snippets used in the experiment. During the adaptive staircase trials, participants were asked to respond with a key press if they heard the sound or not. Some trials contained no stimuli to ensure the participants performed the task according to their hearing capabilities. If participants detected the sound, the volume level decreased (step-down), otherwise the volume level increased (step-up) for the subsequent trial, according to the current step size. When a step-up occurs after a step-down or vice versa, this is called a reversal. At each reversal, the step size decreased until the stopping criterion of 5 reversals was reached. Step sizes of 16, 8, 4, 2, 1 dB were used for the change in loudness before each of the five reversals, respectively. The individual hearing threshold was estimated as the average loudness during the last four reversals. At the end of the adaptive staircase, an example audio snippet (not reused during the main experiment) was played to verify that the final volume was at a comfortable level, and participants had the opportunity to adjust the volume level in 1 dB increments if they perceived it as too loud or too soft, respectively.

### Neurofeedback Training

The neurofeedback training was conducted in two types of experimental blocks: Feedback blocks and Baseline blocks. Each participant completed one (“pre-training”) Baseline block, three Feedback blocks, and another (“post-training”) Baseline block, in that order. Baseline blocks were 21.33 minutes long (40 trials), and Feedback blocks were 16.33 minutes long (28 trials).

In each trial, irrespective of block type, participants were presented with 22-s excerpts from the two audiobooks (see Stimulus materials) and were asked to focus their attention on one of the audiobooks. The order of excerpts was randomised, i.e. it did not follow the logical order from the books.

During Feedback blocks, excerpts from both audiobooks were always presented simultaneously. During Baseline blocks, only one audiobook was presented in 50% of the trials (selected pseudo-randomly). At the start of each trial, the participant was instructed on screen about which audiobook to attend to (female or male speaker). During the sounds, they were asked to fixate on a cross in the middle of the screen. Each block was split into half, with one audiobook as the target in the first half (female or male speaker), and the other in the second half. The target audiobook for the first half of the first experimental block was selected randomly for each participant, followed by alternating targets in the rest of the experiment.

After each trial, regardless of block type, participants were asked to select one out of four possible four-word phrases that were part of the preceding target speech snippet. The four options provided were all part of the target audiobook, but only one of them was part of the preceding snippet. The phrases consisted of four consecutive words but were not necessarily linguistically meaningful in the presented four-word form.

During Feedback blocks only, immediately after the comprehension question, participants were presented with feedback on screen for 3 s, in the form of two bars. In the NF group, the two bars displayed the N1 values from the TRF estimated from responses to target and distractor speech in the preceding 22-s EEG response, respectively, and converted to a percentile as described below (“Online Data Processing”). In the SF group, these values were replayed from another participant from the NF group. The SF group was designed to ensure that any training effects are specific to NF and not driven by other factors like motivation (Alino, 2016; Schabus et al., 2017). Participants from both groups received instructions to increase the “target bar” and lower the “distractor bar”, and were informed that this could be achieved by paying attention to the target speech and ignoring the distracting speech, respectively. They were asked to keep their eyes open and to refrain from moving and tightening their jaw during the task.

### Data Collection and Analysis

Two computers running MATLAB 2019a were used for this experiment (**Fig. 1B**), with one responsible for presenting the stimuli (Presentation computer) and recording the participant’s behavioural response, and the other for processing and recording the incoming EEG data (Backend computer). The Backend computer was used for estimating TRFs and the subsequent neurofeedback calculations, and for sending them to the Presentation computer using Lab Streaming Layer (LSL) (Kothe et al., 2025). Both computers had their experimental states synchronised throughout the experiment using separate LSL streams to prepare the required data for subsequent trials and to communicate and cross-validate their current status at any point during the experiment.

### EEG

EEG was recorded from 64 active electrodes, placed according to the international 10-10 system, using a BioSemi Active two amplifier (BioSemi, Amsterdam, Netherlands) at a sampling rate of 2048 Hz. The electrodes were mounted on an elastic BioSemi head cap and connected to the participant’s scalp via a conductive saline gel (Signagel, Parker Laboratories Inc., Fairfield, NJ, USA). The signal offsets of all electrodes were maintained below 50 µV. The incoming EEG data from the amplifier was received via USB on one of the computers using the BioSemi LSL app (Github) at its native sampling rate of 2048 Hz. EEG recording and pre-processing were performed during the experiment using custom MATLAB scripts and EEGLAB (Delorme & Makeig, 2004) on the same computer, utilising the EEG data stream from the BioSemi LSL app.

### Temporal Response Function (TRF) fitting

All TRFs were calculated using the mTRF-Toolbox v2.3 (Crosse et al., 2016). A constant λ ridge regularisation parameter of 10 was used, based on previous work (Fiedler et al., 2017, 2019; Panela et al., 2024; Yasmin et al., 2023). The constant λ value was favoured over the use of cross-validated optimisation to reduce computational time, given the real-time nature of the experiment. The derivative of the amplitude envelope of our speech stimuli was used as the predictor in the TRF model. The speech envelope and its derivative were extracted individually for each audiobook snippet using a custom MATLAB script. From each snippet, the overall envelope was extracted from 32 different Greenwood-spaced frequency bands after half-wave rectification. The envelope was then down-sampled to 256 Hz, and a derivative was extracted; no further processing was performed on the predictor.

We did not half-wave rectify the envelope derivative and consequently, modelled onsets and offsets with opposite polarities. While neural responses can cancel out in such a model if they are of similar polarity for both onsets and offsets, removing offsets from the predictor through half-wave rectification reduces the amount of data available in the model and might likewise reduce sensitivity in estimating neural responses to speech. We contrasted TRFs from models with and without half-wave rectification of the envelope derivative in our data and show the results in Fig. S1. We found that the attentional modulation of the TRF’s N1 component (target vs distracting speech) is stronger in the model without (Fig. S1A) than with half-wave rectification (Fig. S1B). As our neurofeedback training strongly relies on this attentional effect, this observation validates the inclusion of offsets in the predictor.

The EEG signal was resampled to 256 Hz to match the derivative of the speech envelope, and TRFs were computed for each EEG channel across time lags of -100 to 500 ms between the two. The TRFs were then averaged across a fronto-central montage (F1, Fz, F2, FC1, FCz, FC2, C1, Cz, C2) where we expected to find the maximal N1 component response based on previous literature (Niitinen et al., 1988). Neural responses to targets and distractors were modelled separately.

### Online Data Processing

After each 22-s snippet of audiobooks, the corresponding EEG data was analysed immediately to extract single-trial TRFs. The analysis was done on a dedicated EEG computer that received data via LSL (**Fig. 1B**). EEG triggers synchronised with the presented sounds ensured that 22-s EEG segments were extracted reliably, and matched with the envelope derivatives of the sounds to compute the TRF for the respective trial. Given the fast processing required for online neurofeedback, no EEG artefact attenuation was performed, and a minimal pre-processing pipeline was used. The pipeline consisted of resampling the data to 256 Hz, bandpass filtering it to 1-30 Hz (pass-band), and average referencing. The filter used was a zero-phase Hamming-windowed sinc FIR filter (pop_eegfiltnew function, EEGLAB version 2024.2). The filter order was automatically estimated based on a transition bandwidth of 1 Hz, resulting in a filter order of 847. Cutoff frequencies represent -6 dB attenuation points. The pre-processed data was then used to compute single-trial TRFs, separately for target and distracting speech. The computation of these TRFs took about 120ms per trial.

Single-trial TRFs were converted to feedback (in the NF group and in Feedback blocks only) as follows. The participant’s individual time range was first estimated for the TRF’s N1 component (**Fig. 1C**). This was achieved by computing the TRF, averaged across all trials collected at this point in the experiment (including only multi-speaker trials from Baseline blocks, and all Feedback trials) and the selected EEG channels (see TRF fitting). From this TRF, all the positive and negative peaks in the TRF were extracted using the “findpeaks” MATLAB function. We then extracted *N_peak_* as the most negative peak within a pre-defined time window of 80-150ms. This “search window” was based on previous literature on N1 time lags in envelope-based TRF models (e.g., Orf et al., 2023). We then extracted the nearest positive peaks (minimum distance of 10ms) that precede and follow *N_peak_* in time and were labeled *P_peak_*1 and *P_peak_*2, respectively. The final time range *N_t_*_1_ - *N_t_*_2_ for the individual N1 component was defined as *N_t_*_1_ = *N_peak_* − 0.25(*N_peak_* − *P_peak_*1) and *N_t_*_2_ = *N_peak_* + 0.25(*P_peak_*2 − *N_peak_*), where *P_peak_*1, *P_peak_*2, *N_t_*_1_, *N_t_*_2_, and *N_peak_* are all values in ms. If no negative peaks were found, *N*_t1_ was defined as 107 ms, and *N_t_*_2_ was defined as 123 ms, based on typical N1 lags as defined in Orf et al. (2023). *N_t_*_1_ and *N_t_*_2_ were re-estimated after each experimental block (Baseline or Feedback), and separately for target and distracting speech as well as for each audiobook. After the Pre-Training Baseline, the experimenter manually verified the quality of the estimated TRF and its N1 range. As participants were randomly assigned to groups by the computer and the experimental design was identical for both groups up to this point, the experimenter remained blinded to the participants’ group label. After this point, no further TRFs or lags were visible to the experimenter. After each Feedback trial, the N1 amplitude was computed as the mean amplitude of the TRF between *N_t_*_1_ and *N_t_*_2_. This amplitude was then expressed as a percentile, relative to the distribution of all other single-trial N1 amplitudes measured for that participant until this point in the experiment (including only multi-speaker trials from Baseline blocks, and all Feedback trials). This percentile (between 0 and 100) was the feedback value displayed on the screen. Percentiles were computed separately for target and distracting speech, as well as for the two audiobooks (male and female speakers).

### Offline Data Processing

Several additional processing steps were implemented offline prior to final statistical analysis.

First, noisy EEG channels were identified as those showing discrepant amplitudes from other channels, using a threshold of 3 Kurtosis, and then interpolated using spherical interpolation (Kang et al., 2015). Importantly, EEG channels that were used for neurofeedback were not considered in this step. The statistical evaluation of neurofeedback success and the underlying TRFs were therefore not affected by this processing step. It was only implemented to achieve clean topographies that were required for cluster-based statistics (described below).

Second, we homogenised N1 time ranges (*N_t_*_1_ - *N_t_*_2_) across participants. During neurofeedback, individual time intervals were used as described above. However, this individualisation led to intervals that are broader for some participants than for others, resulting in strong variation in N1 amplitudes. This between-subject variability was not relevant during neurofeedback, as it was given on the individual level. However, it can impact the group-level analysis, as N1 amplitudes are not necessarily comparable across participants as long as individual time ranges are used. Using a common time range for the statistical analysis avoids such a bias. We chose a narrower range than the one used for feedback, as the hypothesised effect of neurofeedback should be maximal at the negative peak of the N1. Due to the availability of the complete dataset for offline analysis, a precise estimate of *N_peak_* and the use of a narrow time range was possible. In contrast, the use of more than a single time point reduced vulnerability to outlier data points. Due to these reasons, the time range for offline data analysis was defined as *N_peak_* ± 8*ms*. We ensured that the use of a broader range does not significantly alter results. *N_peak_* was obtained as described above (Section Online Data Processing), but extracted from the mean TRF across all participants. We verified that *N_peak_* did not differ between audiobooks (male/female) and therefore averaged data across the two conditions before computing *N_peak_*. However, *N_peak_* differed slightly between NF and SF groups and between target and distracting speech (**Fig. 2C**) and was therefore computed separately for these groups and conditions. We verified that the use of a common *N_peak_* for the two groups did not change the study outcomes.

**Figure 2.**
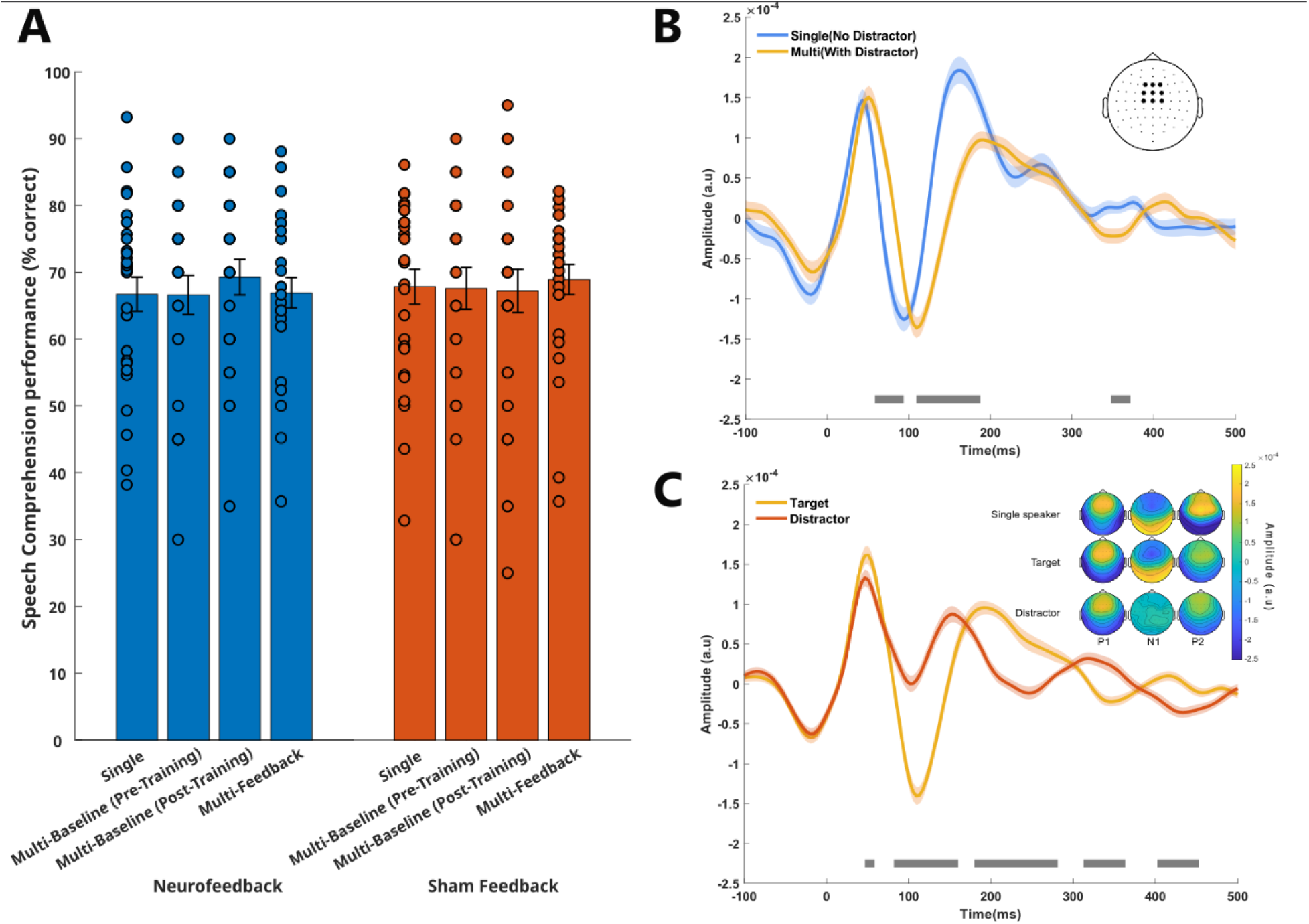
Overview of speech comprehension and TRF results. **A.** Performance in the speech comprehension task. Circles show results from individual participants, bars show the group average, and error bars the standard error of the mean (SEM). **B,C.** TRFs estimated from responses to the derivative of the speech envelope at the EEG channels used for neurofeedback (inset). **B.** TRFs in the absence (“single”) vs presence (“multi”) of a distractor in Baseline blocks. **C.** TRFs estimated for target and distracting speech (Feedback trials and “multi” trials from Baseline blocks). Shaded areas indicate the standard error of the mean (SEM) and grey lines highlight the timepoints where the TRFs are significantly different from each other. Topographies of the most prominent components (P1, N1, P2) from the TRFs are shown as insets and were extracted individually for each condition and component (Single Speaker [P1-39 – 47ms, N1-89 – 97ms, P2-156 – 164ms] from B, Target [P1-46 -54ms, N1- 105 – 113ms, P2- 179 – 187ms] and Distractor [P1- 42 – 50ms, N1- 97 – 105ms, P2- 148 – 156ms] from C).

Third, to rule out the possibility that our results were driven by artefacts in the data, we performed independent component analysis (ICA; Extended Infomax algorithm, EEGLAB) and rejected components automatically classified (with >90% probability) as originating from muscle, eye, cardiac, line noise, or channel noise. We then compared the results with those obtained from data processed using an otherwise identical pre-processing pipeline but without ICA-based artefact correction. The results were essentially unchanged, indicating that our findings were robust to artefact correction. We only report the results obtained from the uncorrected data, as ICA may alter the signals at the channels used for neurofeedback. This approach ensures that the reported responses most closely reflect the signals used to calculate neurofeedback online.

### Statistical Analyses

Statistical analyses were performed in MATLAB along with cluster-based permutation tests (at least two neighbouring channels, 10,000 permutations) from Fieldtrip where appropriate (Oostenveld et al., 2011).

For most analyses, our main factors were Group (NF group vs SF group) and Type (Baseline vs Feedback blocks). In Baseline blocks only, we also contrasted “single-speaker” (only target present) with “multi-speaker” (target and distractor present) trials and pre-training with post-training performance. For all comparisons between Baseline and Feedback blocks, only multi-speaker trials were included in the analyses.

On the behavioural level (**Fig. 2A**), we computed the percentage of correct responses for each participant and condition. We then applied three mixed ANOVAs: one to test for effects of group (between subjects) and block type (within subjects), another for effects of group (between subjects) and training (pre-training vs post-training) in Baseline blocks, and the third for effects of group (between subjects) and distraction (within subjects) in Baseline blocks.

On the neural (EEG) level, we first contrasted TRFs between conditions at each of the lags computed (**Fig. 2B**). For this purpose, we used paired t-tests and used false discovery rate (FDR, Benjamini-Hochberg procedure) correction to account for multiple comparisons.

To test whether neurofeedback modulates the TRF’s N1 component (**Fig. 3**), we designed a linear mixed-effects model (“fitlme” function, MATLAB) of N1 amplitudes with fixed effects “group” and “block type”, and “subject” as a random intercept. The model was computed separately for target and distracting speech, and for each of the 64 EEG channels, and yielded t-values for each main factor and their interaction. These t-values were subjected to cluster-based permutation tests. For our main hypothesis (increased N1 during feedback in the neurofeedback group), we focused the analysis and cluster correction on the 9 EEG channels used to compute feedback values during the training (**Fig. 3A**).

**Figure 3.**
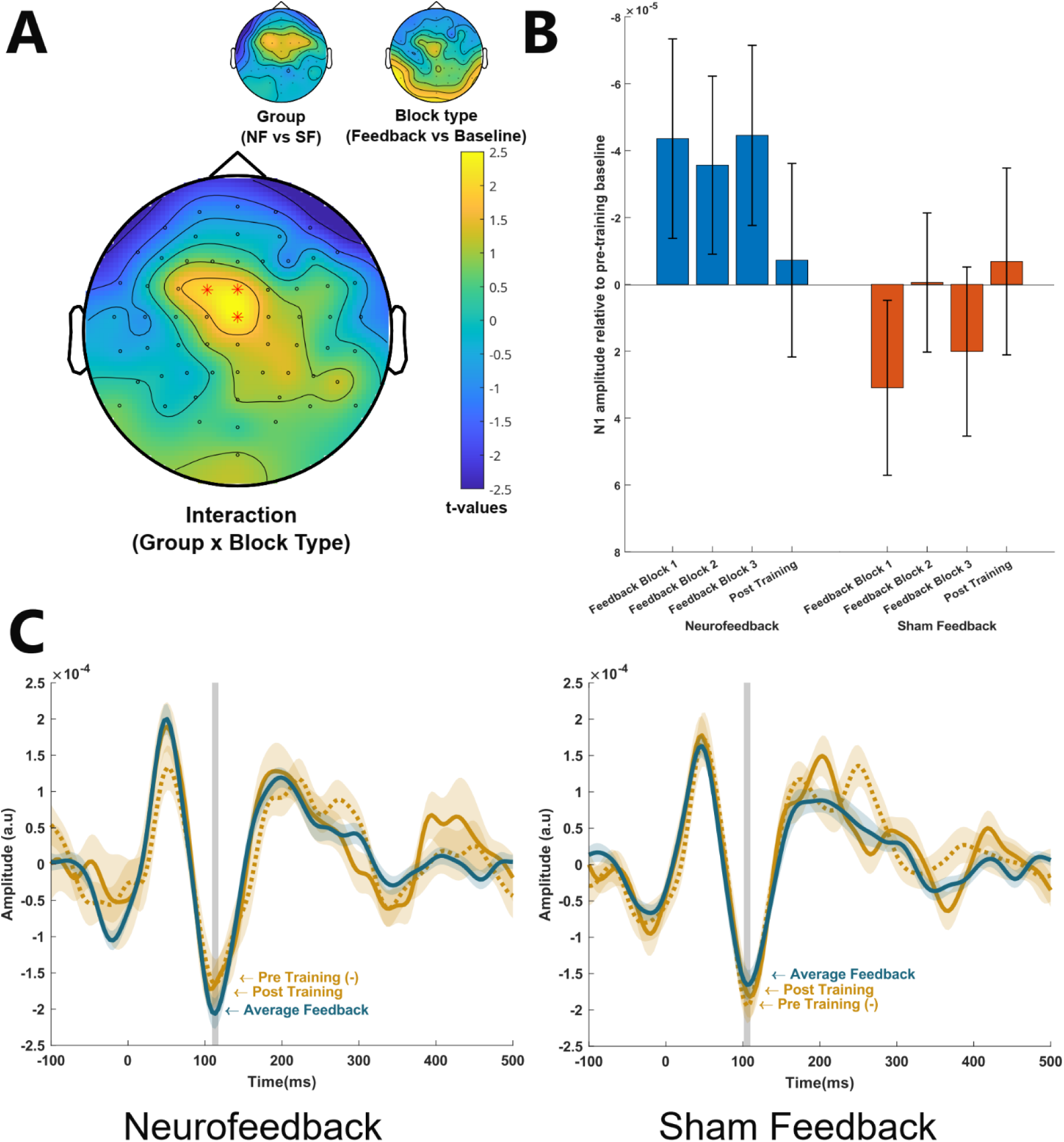
Neurofeedback increases the N1 component in the EEG response to target speech. **A**.T-values from main effects of group (neurofeedback vs sham feedback) and block type (feedback vs baseline) on N1 amplitudes, as well as their interaction in a linear mixed model. The interaction is significant in a cluster comprising the three EEG channels shown by asterisks. A positive interaction term indicates that the increase in N1 negativity from baseline to feedback blocks is greater in the NF group than in the SF group. **B**. N1 amplitude values averaged across the three channels in the significant cluster (Panel A) relative to the pre-training baseline (subtraction within participants). Error bars indicate the SEM. **C**. TRFs estimated from channel FCz, averaged across the three Feedback blocks. Shaded areas around the mean indicate the SEM. Grey lines illustrate the time window used for statistical analysis.

We designed two complementary analyses to test how neurofeedback effects change over time. First, we tested whether effects change during training by fitting a regression line to N1 amplitudes measured in the three Feedback blocks (**Fig. 3B**) and comparing its slope with 0 (t-tests). Second, we tested whether effects persist after training by contrasting N1 amplitudes between post-training and pre-training Baselines (**Fig. 4**). For this purpose, we used a linear mixed-effects model as described above, but replaced the factor “block type” with “time” (pre-training vs post-training). This analysis was done for the EEG channels that exhibited neurofeedback effects (**Fig. 3A**) as well as for the complete 64-channel montage (cluster correction was adapted accordingly).

**Figure 4.**
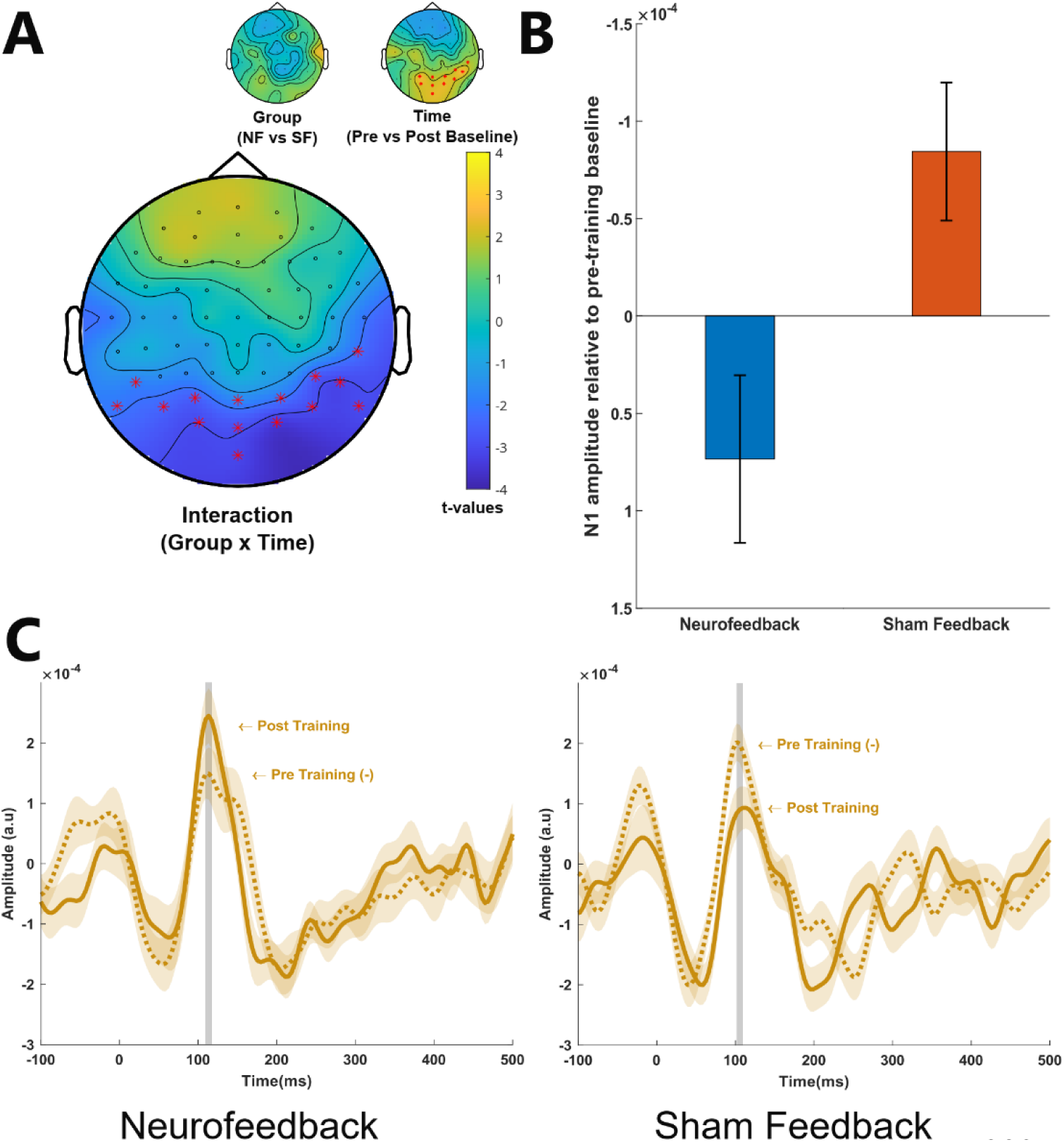
Increased N1 amplitudes in the EEG response to target speech after neurofeedback (during Baseline blocks). **A**.T-values from main effects of group (NF vs SF) and time (pre-training vs post-training) on N1 amplitudes as well as their interaction in a linear mixed model. This interaction is significant in a cluster comprising the fifteen EEG channels shown by asterisks. A positive interaction term indicates that the increase in N1 negativity from pre-training to post-training baseline is greater in the NF group than in the SF group. **B**. Difference (post-minus pre-training Baseline) in N1 amplitude values, averaged across the fifteen channels in the significant cluster (Panel A). Error bars indicate the SEM. **C**. TRFs estimated from channel Iz. Shaded areas around the mean indicate the SEM. Grey lines illustrate the time window used for statistical analysis.

To test whether neurofeedback effects correlate with speech comprehension (“transfer effects”), both N1 amplitudes and performance were expressed relative to those measured in the pre-training baseline for individual participants. This was done as we expected relative (change in TRF or comprehension relative to an individual baseline) rather than absolute effects. We then correlated both relative changes (N1 and comprehension) with each other (Pearson’s correlation). The resulting regression coefficients were Fisher z-transformed and contrasted between NF and SF groups using t-tests (**Fig. 5**). We again performed the analysis (“ft_statfun_indepsamplesregrT” function from Fieldtrip) for the EEG channels that exhibited neurofeedback effects (Fig. 3A) as well as for the complete 64-channel montage (cluster correction was adapted accordingly). We verified results using Shepherd’s pi (Schwarzkopf et al., 2012), a robust correlation procedure that accounts for outliers (Mahalanobis distance m >= 6).

**Figure 5.**
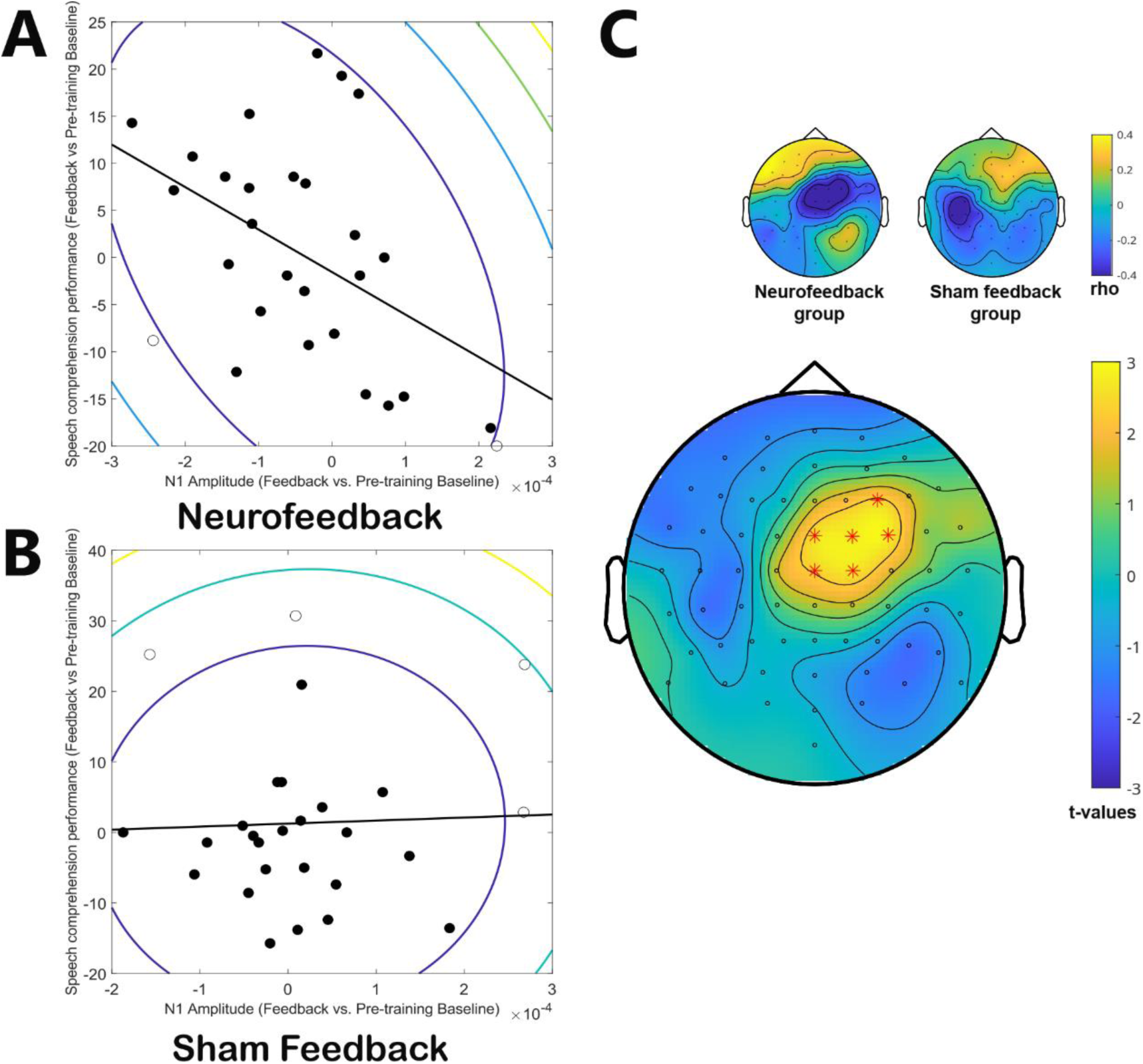
Only in the NF group, increased N1 amplitudes during feedback are associated with better speech comprehension. A,B. Correlation between the average N1 amplitude and speech comprehension performance during Feedback blocks (both relative to pre-training Baseline) in the NF group (A) and SF group (B). N1 amplitudes were computed in the three-channel cluster that showed reliable neurofeedback effects (Fig. 3A). The contour lines indicate the bootstrapped Mahalanobis distance from the bivariate mean in steps of six squared units (Brighter colours denote greater distances). Filled circles denote data included in Shepard’s Pi correlations, open circles denote outliers. The solid line is a linear regression over the data after outlier removal. **C.** The small topographies show correlation coefficients, computed equivalently to panels A,B, but for all 64 EEG channels. The large topography shows t-values from the contrast in correlation coefficients between neurofeedback and sham feedback groups. The contrast is significant in a cluster comprising the six EEG channels shown by asterisks. Positive T-values indicate that a negative correlation between N1 amplitude and speech comprehension performance is stronger in the NF group than in the SF group.

We performed a control analysis (**Fig. 6A,B**) to ensure that N1 values are indeed reliable on the single-trial level, as assumed by our neurofeedback training. For this purpose, a sensitivity index was calculated for each participant, and separately for target and distractor processing. It was calculated by dividing the average of single-trial N1 amplitudes (multiplied by -1), from the channels used for feedback, by their standard deviation. To correct for effects of baseline differences (e.g., most TRF values larger or smaller than 0), single-trial N1 amplitudes were expressed relative to the transition point (50%) between P1 and N1 of the same trial. The resulting values from single participants were contrasted with 0 (t-test) to obtain a measure of reliability on the group level. We also calculated the difference between the feedback values presented to the sham feedback group (**Fig. 6C**) by contrasting the difference between the value they received to the value that would have received during genuine neurofeedback (as percentiles)

**Figure 6.**
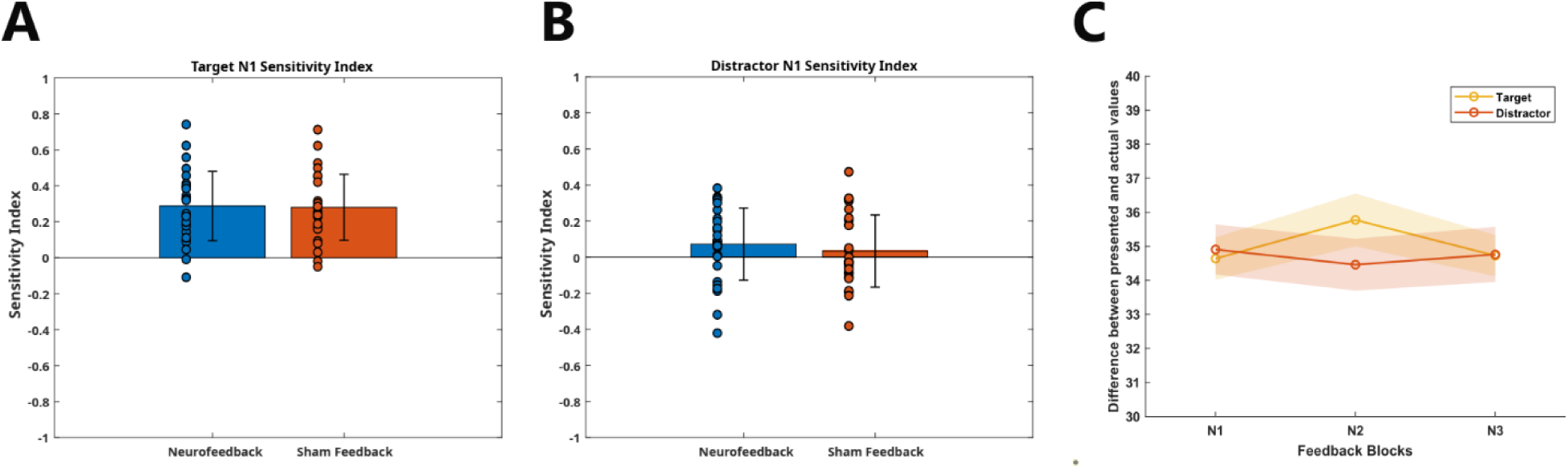
Control analyses. A,B. Reliability of single-trial N1 values. Single-trial N1 amplitudes from TRFs (multiplied by -1 and relative to a baseline) were estimated for target (A) and distracting speech (B), and their average was divided by their trial-to-trial variability. Circles show outcomes from individual participants, bars and error bars show the mean and SEM across participants, respectively. **C. Feedback vs actual N1 values in the sham feedback group.** Average distance between displayed feedback values (as percentiles) in the SF group (C) and those that would have been displayed during genuine neurofeedback. Shaded areas represent the SEM.

Finally, we explored whether neurofeedback modulates the TRF’s P1 component (**Fig. 7**). Similar to our analysis with the N1, we designed a linear mixed-effects model (“fitlme” function, MATLAB) of Target speech P1 amplitudes with fixed effects “group” and “block type”, and “subject” as a random intercept and for each of the 64 EEG channels, and yielded t-values for each main factor and their interaction. These t-values were subjected to cluster-based permutation tests, first on the 9 EEG channels used to compute feedback values during the training and subsequently with all 64 channels. To test whether the neurofeedback effects were linked, we examined whether any potential change in P1 was associated with changes in N1 by comparing Pearson’s correlations between their amplitudes across all channels within their respective significant clusters.

**Figure 7.**
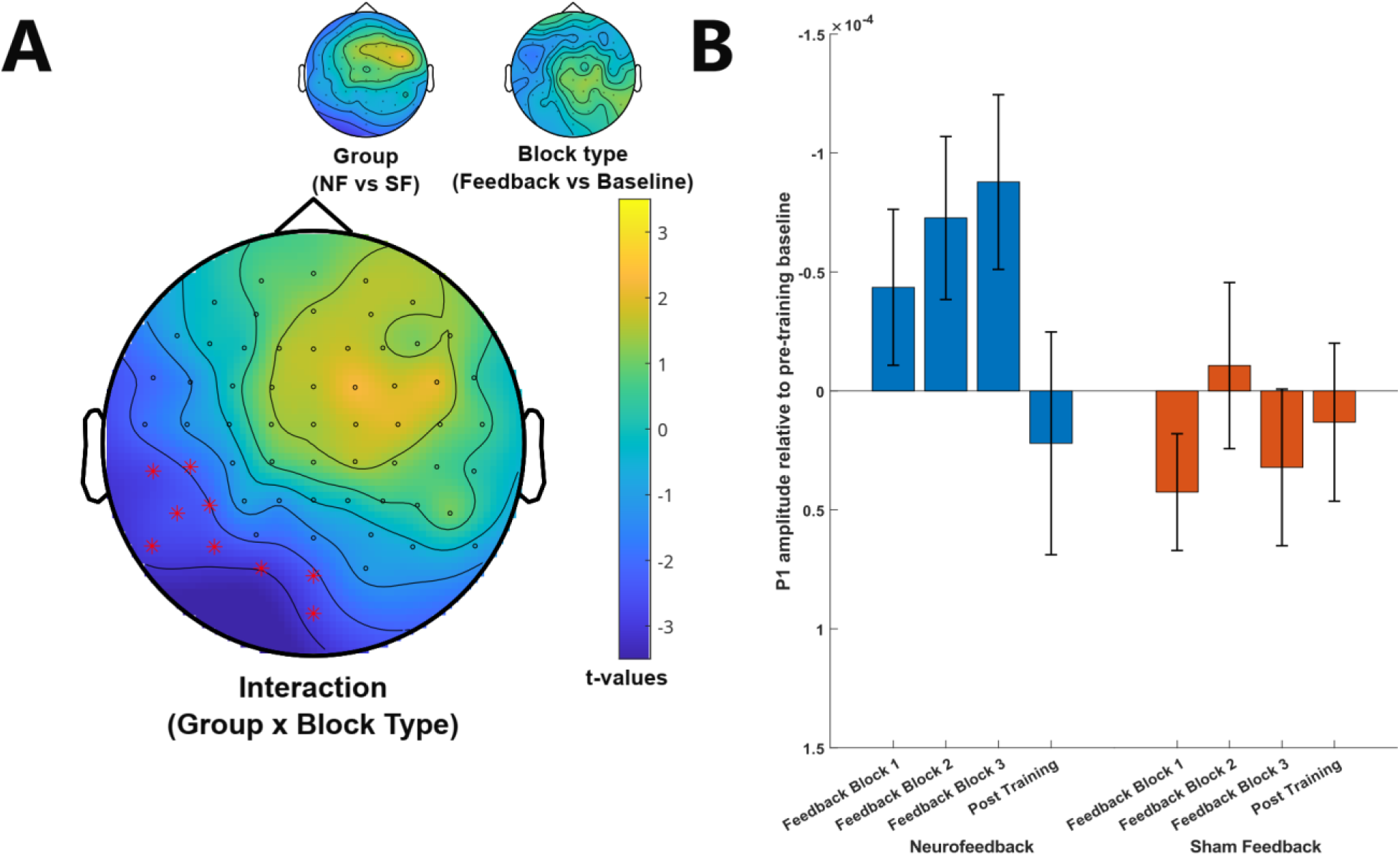
Increase in P1 amplitude during Feedback only for the neurofeedback group. **A.** T-values from main effects of group (NF vs SF) and block type (feedback vs baseline) on P1 amplitudes, as well as their interaction in a linear mixed model. This interaction is significant in a cluster comprising the nine EEG channels shown by asterisks. A positive interaction term indicates that the increase in P1 positivity from baseline to feedback blocks is greater in the NF group than in the SF group. **B**. P1 amplitude values averaged across the nine channels in the significant cluster (Panel A) relative to the pre-training baseline (subtraction within participants). Error bars indicate the SEM.

## Results

### Overview

Participants completed three Feedback (“training”) blocks, preceded by one Baseline block (“pre-training”) and followed by another one (“post-training”) (**Fig. 1A**). Hereafter, the term “Baseline block” refers to both the pre-training and post-training blocks, unless specified otherwise. In all blocks, participants attended to one of the two concurrently presented audiobooks. One trial consisted of a 22-s segment of speech, followed by a comprehension question (see Study Design). In Baseline blocks, no distractor was present in half of the trials. In Feedback blocks only, the comprehension question was followed by a visual display of the participant’s N1 amplitudes (**Fig. 1A**). These amplitudes were extracted from TRFs estimated from responses to the derivative of speech envelopes at fronto-central EEG channels during the preceding speech segment (**Fig. 1B,C**). Feedback was given separately but simultaneously for responses to target and distracting speech. In the NF group (N=28), N1 amplitudes were expressed as a percentile (between 0 and 100) relative to all other N1 amplitudes measured up to that point in the experiment (see Online Data Processing). Participants were aware that these values reflect their attentional focus. They were told to control them by paying attention to the target audiobook and ignoring the distractor, respectively. In the SF group (N=27), each participant was presented with a replay of the feedback given to a participant in the NF group, but received the same instructions.

We expected that successful neurofeedback training would strengthen the N1 component in responses to target speech (relative to Baseline blocks) and weaken the N1 in responses to distracting speech, but only in the NF group. Such an effect would be visible as an interaction between block type (Feedback or Baseline) and group (NF or SF) on N1 amplitudes in a linear mixed model (see Statistical Analysis). While this analysis does not distinguish pre- and post-training Baselines, it is possible that neurofeedback effects persist after training (e.g., stronger N1 in the post-training Baseline than in the pre-training Baseline) or develop during training (e.g., N1 that increases over time). These effects were explored in follow-up analyses.

### Speech comprehension is stable across groups and conditions

In each trial, participants selected one out of four four-word phrases that they believed was part of the preceding target speech. **Fig. 2A** shows performance in this task and how it varied with group (NF or SF), block type (Baseline or Feedback), and the presence of a distractor (note that single-speaker trials were tested in Baseline blocks only).

In all conditions and groups, participants performed well above the chance level of 25 %. However, no statistical differences were observed between the two groups (*F* = 0.05, *p* = .821), nor between block types (*F* = 0.04, *p* = .83), or the interaction of group and block type (*F* = 1.20, *p* = .278). While, only in the Neurofeedback group, post-training performance in Baseline blocks was numerically better than pre-training performance (**Fig. 2A**), this difference was not significant (pre- vs post-training Baseline: F = 0.27, p = .608; interaction with group: F = 0.46, p = .499). Moreover, the addition of a distractor did not reduce performance as expected (*F* = 0.16, *p* = .694; “single” vs “multi” in Baseline blocks only). Although these results seem to reveal neither distraction effects nor neurofeedback effects on speech comprehension, such effects did emerge in more detailed neural analyses as reported below.

### The N1 component of the Temporal Response Function is a sensitive measure of attention

**Figs. 2B,C** give an overview of the TRFs obtained across all participants (N=55) and the nine EEG channels used for neurofeedback (inset in **Fig. 2B**). Horizontal lines illustrate time points with significant differences between TRFs shown (FDR-corrected p < 0.05 from paired t-tests).

**Fig. 2B** shows the TRF estimated for target speech in Baseline blocks, when only one audiobook was presented (blue line) and how it changed with the addition of a distractor (yellow line). Not only did the distractor delay the N1 to target speech, it also reduced later response components (between 100 ms and 200 ms). This finding shows that, despite stable measures of speech comprehension (**Fig. 2A**), the addition of a second speech stream did change the neural responses estimated for target speech, possibly through the increased cognitive load for participants.

**Fig. 2C** contrasts TRFs estimated from target and distracting speech, respectively, averaged across all blocks. As expected, we found that attention strongly increased some TRF components, particularly the N1 (around 100 ms). This finding supports the notion that the N1 is indeed a sensitive measure of attentional focus, supporting a possible efficacy of our neurofeedback training.

### Neurofeedback increases the N1 component in the response to attended but not distracting speech

Having established the N1 component of the TRF as a sensitive marker of attention to speech, we then tested our main hypothesis that it can be trained with neurofeedback. A linear mixed-effects model was used to test whether block type (Feedback or Baseline, within-subjects) and group (NF or SF, between-subjects) modulated the N1 amplitude. The model was computed separately for neurofeedback effects on target and distractor processing, and for each of the 64 EEG channels. For our main hypothesis (increased N1 during feedback in the neurofeedback group), we focused our statistical analysis on the 9 EEG channels used to compute feedback values during the training. While individual time lags for the N1 component were used for neurofeedback, we extracted common lags (NF group: 109 – 117 ms, SF group: 101 – 109 ms; grey window in **Fig. 3C**) for the group-level analysis (see Statistical Analyses for the underlying rationale).

**Fig. 3** shows outcomes from this analysis for target speech, averaged across both audiobooks. Results revealed marginally increased N1 amplitudes in the NF group (group effect in **Fig. 3A**; max. *t* = 1.81, min. *p* = 0.071 across the 9 feedback channels) but not during Feedback blocks (block type effect in **Fig. 3A**; max. *t* = 1.36, min. *p* = .175). None of these main effects survived cluster correction. However, we found a statistically significant interaction between block type and group in a cluster that comprised the channels FCz, Cz, and FC1 (Cluster *t* = 6.80, *p* = .014) (**Fig. 3A**). Post-hoc t-tests revealed larger N1 amplitudes in Feedback blocks than in Baseline blocks, but only in the NF group (*t* = 2.36, *p* = .013), and not in the SF group (*t* = -1.22, *p* = .883). **Fig. 3B** illustrates this finding by showing N1 amplitudes relative to the pre-training baseline (subtraction within subjects). N1 amplitudes exceed pre-training levels in all Feedback blocks in the NF group but in none of them in the SF group, an observation we come back to later. **Fig. 3C** illustrates the underlying TRFs from a central channel in the cluster (FCz). An increased N1 during Feedback blocks as compared to both baselines is again visible in the NF group, whereas such an effect is not present in the SF group. Together, we confirmed that the N1 is a neural signature of attention to speech, and found that it can be enhanced through neurofeedback training. Further support for the efficacy of the training comes from the finding that neurofeedback effects were specific for a subset of EEG channels that were used for the feedback (cf. topographies in **Fig. 3A**).

We then performed the same statistical tests on neural responses to distracting speech. As shown in **Fig. S2**, we did not find significant main effects of group (max. *t* = 0.21, min. *p* = .813 across the 9 feedback channels) or block type (max. *t* = 1.88, min. *p* = .062), nor an interaction between the two (max. *t* = 0.4780, min. *p* = .633). One potential reason for this absence of neurofeedback effects on distractor processing is that corresponding neural responses are comparably small (see Results section: Reliability of single-trial N1 values).

### Persistent effects on the N1 component in the neural response to attended speech after neurofeedback

Having established the effects of neurofeedback on the N1 component of TRFs estimated for target speech, we tested how these effects develop over time. A regression line was first fitted to the N1 amplitude (averaged across the three channels in the significant cluster; **Fig. 3A**) across the three Feedback blocks (cf. **Fig. 3B**) to test whether this amplitude changes linearly over time. In neither of the two groups, the slope of this line differed significantly from 0 (NF group: *t* = -0.04, *p* = .967 / SF group: *t* = -0.44, *p* = .662). The slope did not differ between groups either (*t* = 0.29, *p* = .771), indicating that neurofeedback effects were consistent and did not increase over time as it is sometimes observed (Enriquez-Geppert et al., 2026; Zoefel et al., 2011).

We then tested whether neurofeedback effects persisted after training by contrasting N1 amplitudes between the two Baseline blocks. We initially expected such persistent effects in the same channels that exhibit neurofeedback effects (**Fig. 3A**). However, it is already visible from **Fig. 3B** that this expectation was not confirmed, and we consequently tested for effects across all 64 EEG channels (with corresponding correction for multiple comparisons). We found an effect of time (pre-training vs post-training) in an 11-channel cluster (Cluster *t* = -24.05, *p* = .051) that was slightly above the conventional significance threshold of α = 0.05 and not investigated further due to an interaction effect in a similar set of channels. As shown in **Fig. 4A**, we found a statistically significant interaction between time (pre- vs post-training) and group (NF vs SF) in a cluster that comprised 15 parieto-occipital EEG channels (Cluster *t* = 38.69, *p* = .016). **Figs. 4B, C** demonstrate that this interaction was driven by an N1 component that became stronger after training in the NF group, but weaker in the SF group (note that the N1 is positive in parieto-occipital channels). Similar effects with opposite polarity are visible in fronto-central EEG channels (**Fig. 4A**), although these did not survive correction for multiple comparisons. Note that this dipolar pattern is often observed for the N1 component in auditory responses, including the current study (see topographies in **Fig. 2**), and does not indicate an involvement of the occipital cortex.

### Increased N1 amplitudes during neurofeedback are associated with improved speech comprehension

We next tested whether a change in N1 amplitude, estimated from responses to target speech, was associated with improved speech comprehension, possibly caused by improved attention to speech. Our results described above revealed training-induced changes in N1 amplitude during all Feedback blocks (**Fig. 3**) and the second (post-training) Baseline block (**Fig. 4**). For each participant, we therefore computed their N1 amplitude in these blocks, relative to their individual pre-training Baseline, and correlated it with the corresponding change in task performance (between-subject correlation). If neural effects on speech comprehension are due to neurofeedback training, these correlations should be stronger in the NF group than in the SF group. We therefore contrasted the resulting correlation coefficients between groups (see Statistical Analyses).

We first focused on the increased N1 amplitudes observed in a three-channel cluster (**Fig. 3A**) during Feedback blocks in the NF group. We found a significant association between increased N1 amplitudes and improved speech comprehension in the NF group (Pearson’s *r* = - .46, *p* = .013) (**Fig. 5A**), but not in the SF group (Pearson’s *r* = .04, *p* = .848) (**Fig. 5B**). Shepard’s Pi correlation analysis confirmed that these results are not driven by outliers (NF group *pi* = - .43, *p* = .054 / SF group *pi* = - .01, *p* = 1). Correlation coefficients differed between the two groups, albeit slightly above the conventional standard significance threshold (*t* = 1.8914, *p* = 0.059). We extended the analysis to all 64 EEG channels and identified a cluster of 6 fronto-central EEG channels where this correlation significantly differed between groups (Cluster *t* = -15.57, *p* = .028) (**Fig. 5C**). Thus, we found that neurofeedback did not only increase N1 amplitudes, for some right lateralised EEG channels these neural changes were also linked to improved speech comprehension.

We did not find any significant correlation between N1 amplitudes and speech comprehension performance in the post-training (relative to pre-training) Baseline block, nor any reliable difference in this correlation between the two groups (*t* = - 0.77, *p* = .442; SF group: Pearson’s *r* = .04, *p* = .851, NF Group: Pearson’s *r* = - .18, *p* = .360).

### Control Analysis: Reliability of single-trial N1 values

We also explored whether single-trial N1 values are reliable enough for meaningful neurofeedback. To do so, we computed, for each participant, the single-trial N1 amplitudes (multiplied by -1 and relative to a baseline; see Statistical Analyses) and divided the resulting average amplitude with their trial-to-trial variability. This comparison is mathematically analogous to signal detection theory’s d′. If its outcome exceeds 0, it suggests that N1 values are not only meaningful in the grand average but also present at the level of individual trials. The outcome highlighted a significant N1 component when estimated from responses to target speech (NF group: *t* = 7.88, *p* = <.001; SF group: *t* = 7.93, *p* = <.001) (**Fig. 6A**) but not for responses to the distractor (NF group: *t* = 1.92, *p* = .066; SF group: *t* = 0.89; *p* = .382) (**Fig. 6B**). This result suggests that N1-based neurofeedback was indeed meaningful for target speech. However, it also suggests that the N1 component in response to distracting speech is, despite its prominent deflection on the average level (**Fig. 2C**), not strong enough to be reliable on the single-trial level. This finding can explain the absence of neurofeedback effects on neural responses to distracting speech.

### Control Analysis: Feedback vs. actual N1 values in the sham feedback group

We verified that the feedback values presented to the SF group were different from their actual N1 values during Feedback blocks. To this end, we computed the distance (absolute difference) between the displayed feedback values and those that the participant would have seen during genuine neurofeedback (both as percentiles). **Fig. 6C** shows this distance, separately for the three Feedback blocks and for the target and distractor-based feedback. We found that the feedback displayed to the SF group diverged from their actual responses by approximately 35 percentiles across all blocks and conditions. This result confirms that the feedback in the SF group was most likely uninformative to them.

### N1 Neurofeedback changes the P1 component in the neural response to attended speech

While our neurofeedback training was tailored to the N1 component of the TRF, results displayed in **Fig. 3C** suggest concomitant effects on the earlier component P1 (∼50 ms). Such effects were unexpected and therefore need to be considered as exploratory. We applied the same linear mixed-effects model to P1 amplitudes as described above for N1 amplitudes (**Fig. 3**). Again, we used common lags (NF group: 46– 54ms, SF group: 42 – 50ms) for group-level analysis. We did not find significant effects of neurofeedback on the P1 component estimated from the 9 channels used for feedback. However, we found a statistically significant interaction (**Fig. 7A**) between block type (Feedback vs Baseline) and group (NF vs SF) in a cluster comprising 9 parieto-occipital EEG channels (Cluster *t* = 23.34, *p* = .031), driven by a stronger P1 component during Feedback blocks, exclusively in the NF group (**Fig. 7B**). Main effects of group (max *t* = 2.22, min *p* = .028) and type (max *t* = -1.69, min *p* = .091) did not survive correction for multiple comparisons.

Given neurofeedback effects on both P1 and N1 components, we tested whether they linked by correlating the two across participants (both from their respective channel clusters). However, we found no reliable correlation between the two (NF group Pearson’s *r* = -.27, *p* = .166 / SF group Pearson’s *r* = -.33, *p* = .091), indicating two independent (rather than concomitant) effects.

## Discussion

In this study, we used neurofeedback to enhance selective attention to speech in a challenging multi-speaker scenario. Neurofeedback training was based on the TRF’s N1 component, a reliable marker of attentional focus as confirmed here (**Fig. 2C**). During feedback, we found an increased N1 component in the response to target speech (**Fig. 3**). Importantly, this increase in a neural marker of attention to speech (1) was only observed during neurofeedback but not sham feedback, (2) was specific for a subset of EEG channels that were used for feedback, and (3) correlated with changes in speech comprehension (**Fig. 5**). Together, these results suggest that neurofeedback can enhance a signature of successful attention to speech, with important implications for individuals that struggle with speech perception in challenging situations (discussed below).

We also obtained results that we had not expected initially. These include post-training effects in parieto-occipital channels (**Fig. 4**), no modulation of distractor processing, and effects on earlier response components (**Fig. 7**). We discuss these results below.

### TRF-based neurofeedback can enhance neural responses to continuous speech

The key result of the present study is that neurofeedback training improved specific components of neural responses to naturalistic speech. This is a remarkable departure from the typical use of neurofeedback, often designed to modulate spectral power in certain frequency bands or responses evoked by simple events (Arvaneh et al., 2019; Enriquez-Geppert et al., 2014; Hayashi et al., 2022; Ruddy et al., 2018; Zoefel et al., 2011). Few studies have attempted neurofeedback training to support neural responses to natural, running speech, most previous work was restricted to isolated words (Kim et al., 2021) or used decoding accuracy for feedback (Haro et al., 2025; Jaeger et al., 2020; Rotaru et al., 2026), rather than a single response component related to selective attention. We here report that neurofeedback training of such specific response components is possible and its effects observed immediately, i.e. in the first training block. Our results open the door to neurofeedback training that involves other dynamic and naturalistic stimuli, such as responses to lip movements that can be quantified using TRFs (Reisinger et al., 2025). An advantage of our TRF-based training protocol is the possibility of providing participants with clear instructions of how their response can be regulated: They were informed that training success can be achieved by focusing attention on the target speech, and away from distractors. During traditional neurofeedback training (e.g., based on oscillatory power), participants often have to find their own strategy to regulate the neural measure of interest, leading to inconsistencies across participants and often to long protocols. Clear instructions may have contributed to the fact that neurofeedback effects were visible from early on during the training.

Results displayed in **Fig. 3C** imply that neurofeedback of the N1 component entails a concomitant regulation of an earlier (P1) component. Yet, neurofeedback effects on the P1 and N1 components affected different EEG channels (**Fig. 3A** vs. **Fig. 7A**) and were not correlated with each other. We therefore conclude that a specific regulation of the N1 component is possible in principle, and effects on the P1 remain to be investigated through future studies.

Contrary to our expectations, we found no neurofeedback effects on neural responses to distracting speech. As participants received simultaneous feedback on their responses to both target and distracting speech, most of them may have chosen to focus on target speech and to ignore distractor feedback. Since previous studies found attentional modulation of distractor speech processing using TRF (Holtze et al., 2021; Straetmans et al., 2022), it seems feasible to train responses to target and distracting speech in different blocks or groups, to test whether such separate feedback makes distractor-based suppression easier. Another explanation is that neural responses to ignored stimuli are inherently smaller and therefore less reliable on the single-trial level. Our control analysis (**Fig. 6B**) supports this assumption. Less reliable neural measures are less informative for feedback and this may have prevented neurofeedback effects on distractor processing. Another point to note is the strong spatial separation in our speaker configuration (target at 0°, distractor at 180°), which differs from the more common lateralized (Left/Right) or co-localized (target and distractor from the same source) layouts (Fiedler et al., 2019). We did not use lateralized configurations as they encourage monaural listening strategies and create lateralized brain activity that is more challenging to interpret. Instead, we used spatially separated (rather than co-localized) speakers to mimic everyday scenarios where targets are often located in front of the listener and distractors are behind them (e.g., in a restaurant). Moreover, our training was designed for future application in clinical use (see below). A strong spatial separation facilitates the training and, consequently, its application in populations that struggle to comprehend speech. Despite its spatial separation from the target, the distractor delayed the N1 component in the response to target speech and reduced later response components associated with cognitive load (**Fig. 2B**) (Alyan et al., 2023; Getzmann et al., 2016). Nevertheless, our choice to spatially separate speakers may have further contributed to a decreased response to distractors. In contrast to our study, Haro et al. (2025) did find reduced responses to distracting speech during TRF-based neurofeedback, which we posit could be due to the use of reconstruction accuracy as their feedback measure. Although reconstruction accuracy is not necessarily driven by attention-related components and might therefore be less efficient for attentional training, it relies on more data and might be more reliable for single trials. It remains to be confirmed whether, for this reason, reconstruction accuracy is more suitable for the training of distractor processing, a hypothesis that can be tested in studies that contrast both approaches (N1 vs reconstruction accuracy) more directly.

### Neurofeedback effects on target processing are linked to speech comprehension

For some EEG channels, an increase in N1 during Feedback blocks was linked to a corresponding enhancement of speech comprehension (**Fig. 5C**). This correlation was only present in the NF group and in a cluster of channels that overlapped with that showing a N1 modulation during neurofeedback (**Fig. 3A**), suggesting that the enhanced comprehension was indeed driven by the training.

At first glance, it therefore seems surprising that we did not find an overall enhancement of speech comprehension in the NF group (**Fig. 2A**). However, the net effect of neurofeedback on speech comprehension should be determined by the *combination* of neurofeedback effects on N1 amplitude (**Fig. 3A**) *and* the link between N1 amplitude and speech comprehension (**Fig. 5C**) in *all* EEG channels. We identified a correlation between N1 amplitude and speech comprehension (stronger N1 amplitude associated with improved comprehension) in a right-lateralised cluster of fronto-central electrodes (**Fig. 5C**). Nevertheless, some left-lateralised channels (e.g., C3, FC1) showed an opposite effect (stronger N1 associated with poorer comprehension). Although this effect did not reach conventional significance, it may have counter-acted the right-lateralised improvements and explains why overall comprehension was not improved in the NF group. Nevertheless, this result yields the concrete prediction that a right-lateralised neurofeedback (using the cluster identified in **Fig. 5C**) would improve speech comprehension, a hypothesis to be tested in future work. In line with our right-lateralised cluster, previous studies have also demonstrated a rightward lateralisation of the N1 component, not only in ERPs (Bardy et al., 2014; Okamoto et al., 2009; Okamoto & Kakigi, 2015), but also in TRFs estimated from neural responses in the presence of competing speech (Ding & Simon, 2012; Mai et al., 2025). This finding is in line with Poeppel’s (2001, 2003) hypothesis that auditory perception is handled asymmetrically by the brain, depending on the relevant timescales in the stimulus. According to the ‘Asymmetric sampling in time’ hypothesis (AST), the left hemisphere uses shorter (∼20-40ms) temporal integration windows than the right hemisphere (∼150-250ms). As we used TRFs estimated for neural responses to the speech envelope and the latter is dominated by timescales between ∼150 and 400 ms (Varnet et al., 2017), it is possible that we measured a neural process that is right-lateralised. Although the AST remains debated, it is supported by some work (see Oderbolz et al., 2025, for an in-depth review) and can explain our observation.

### Outlook

While neurofeedback studies are often conducted in multiple sessions, we found neural and behavioural training effects in a single session. The efficacy of the training may have benefitted from our focus on a neural measure sensitive to attention, and from defining clear regulation strategies in the instructions to participants (see above). The use of multiple sessions in future work may also help to address open questions. For example, our main results demonstrate a successful modulation of neural responses during training, but might not reflect learning processes necessary for longer-term improvements. The persistent neural effects of neurofeedback training we found were relatively weak in channels used for feedback calculation, but stronger in others (**Fig. 4A**). Moreover, these effects were not linked to changes in speech comprehension. It is possible that persistent learning effects require multiple sessions. Achieving such long-term training effects is a key aim for the future, in particular in view of potential clinical applications.

Another open factor is the use of the N1 component as our neural marker of attention. The N1 component is often highlighted as a neural signature of selective attention and is reduced in response to ignored or distracting speech (Orf et al., 2023). Our results confirm this assumption (**Fig. 2C**); however, attention to speech may also involve processes in the auditory pathway that are reflected by other response components (Broderick et al., 2018; Gillis et al., 2023; Karunathilake et al., 2025). Future studies could explore the use of more complex measures for neurofeedback training, such as the N1-P1 complex or the N1-P2 complex, measures that are sometimes used to quantify capture attentional processes (Mai et al., 2025). They could also use different regressors to target the processing of specific parts of speech, such as phonemes, or even visual cues such as lip movements (Reisinger et al., 2025; Teoh et al., 2022).

It remains an open question whether neurofeedback can be used to systematically train distractor suppression. As discussed above, TRF-based reconstruction accuracy may be an appropriate neural measure for this purpose (Haro et al., 2025). However, when using reconstruction accuracy as a neurofeedback signal, Rotaru et al. (2026) found that acoustic feedback can increase distraction and hinder, rather than facilitate, neural and behavioural responses to speech. Notably, feedback in their study was continuously delivered, during the speech stimulus, and updated every second, rather than being delayed until after the stimulus, as in the present work. This finding suggests that both the choice of neural target and the temporal dynamics of feedback delivery jointly shape whether neurofeedback facilitates selective listening. Moreover, other possibilities exist and need to be explored. For example, neurofeedback protocols could be designed that target later components of the TRF (e.g., the P2), which may be functional in active distractor suppression (Fiedler et al., 2019). The efficacy of such protocols remains unknown and testing them is an exciting endeavour for the future.

### Clinical Implications

The ability to understand speech in noise declines with increasing age. An important aspect here is the reduced ability to suppress distractors rather than an actual hearing loss (Füllgrabe et al., 2015; Gillingham et al., 2018; Tun et al., 2009). This suggests deficits at central stages of auditory processing rather than purely peripheral impairment. Improving speech in noise perception can be challenging with conventional methods like hearing aids, and alternative methods remain to be established. Our results suggest that TRF-based neurofeedback may become such an alternative method. Studies with multiple training sessions in populations that struggle with speech-in-noise intelligibility are needed to investigate whether this approach is indeed promising. A complementary or simultaneous application of neurofeedback with existing behavioural training (Burk et al., 2006; Fisher et al., 2025; Sweetow & Sabes, 2006) is a plausible scenario that merits further research.

## Conclusion

We here highlight a novel neurofeedback approach to modulate a neural signature of attention to speech in noise. We demonstrate that a specific, attention-related component of the TRF, estimated from the EEG response to target speech, can be trained successfully even in a single session. Training success is linked to speech comprehension, but lateralisation and persistent effects require further research to be fully understood.

## Data and Code Availability

Data analysis code is available in the following repository: https://doi.org/10.5281/zenodo.20446612

Data will be made available upon publication in the same repository.

## Acknowledgments

This study has been supported by the Fondation pour l’Audition (grant number FPA-RD-2021-10) and through the grant EUR CARe N°ANR-18-EURE-0003 in the framework of the Programme des Investissements d’Avenir.

## Supplementary materials

**Consensus on the Reporting and Experimental Design of clinical and cognitive-behavioural Neurofeedback studies (CRED-nf) best practices checklist 2020**

| Domain | Item # | Checklist item | Reported on page # |
|---|---|---|---|
| **Pre-experiment** |  |  |  |
|  | 1a | Pre-register experimental protocol and planned analyses | NA |
|  | 1b | Justify sample size | Pg. 3 |
| **Control groups** |  |  |  |
|  | 2a | Employ control group(s) or control condition(s) | Sham Feedback group (Pg. 6) |
|  | 2b | When leveraging experimental designs where a double-blind is possible, use a double-blind | Pg. 3 |
|  | 2c | Blind those who rate the outcomes, and when possible, the statisticians involved | NA |
|  | 2d | Examine to what extent participants and experimenters remain blinded | NA |
|  | 2e | In clinical efficacy studies, employ a standard-of-care intervention group as a benchmark for improvement | NA |
| **Control measures** |  |  |  |
|  | 3a | Collect data on psychosocial factors | NA |
|  | 3b | Report whether participants were provided with a strategy | Pg. 5, 22 |
|  | 3c | Report the strategies participants used | NA |
|  | 3d | Report methods used for online-data processing and artifact correction | Pg.7 |
|  | 3e | Report condition and group effects for artifacts | NA |
| **Feedback specifications** |  |  |  |
|  | 4a | Report how the online-feature extraction was defined | Pg.8 |
|  | 4b | Report and justify the reinforcement schedule | Pg.6 |
|  | 4c | Report the feedback modality and content | Pg.5,6 |
|  | 4d | Collect and report all brain activity variable(s) and/or contrasts used for feedback, as displayed to experimental participants | Pg.3,5 |
|  | 4e | Report the hardware and software used | Pg.6 |
| **Outcome measures** |  |  |  |
| Brain | 5a | Report neurofeedback regulation success based on the feedback signal | Pg. 13 |
|  | 5b | Plot within-session and between-session regulation blocks of feedback variable(s), as well as pre-to-post resting baselines or contrasts | Pg. 17 |
|  | 5c | Statistically compare the experimental condition/group to the control condition(s)/group(s) (not only each group to baseline measures) | Pg. 13 |
| Behaviour | 6a | Include measures of clinical or behavioural significance, defined a priori, and describe whether they were reached | Pg. 11 |
|  | 6b | Run correlational analyses between regulation success and behavioural outcomes | Pg. 18 |
| **Data storage** |  |  |  |
|  | 7a | Upload all materials, analysis scripts, code, and raw data used for analyses, as well as final values, to an open access data repository, when feasible | NA |

---

**Figure S1.**
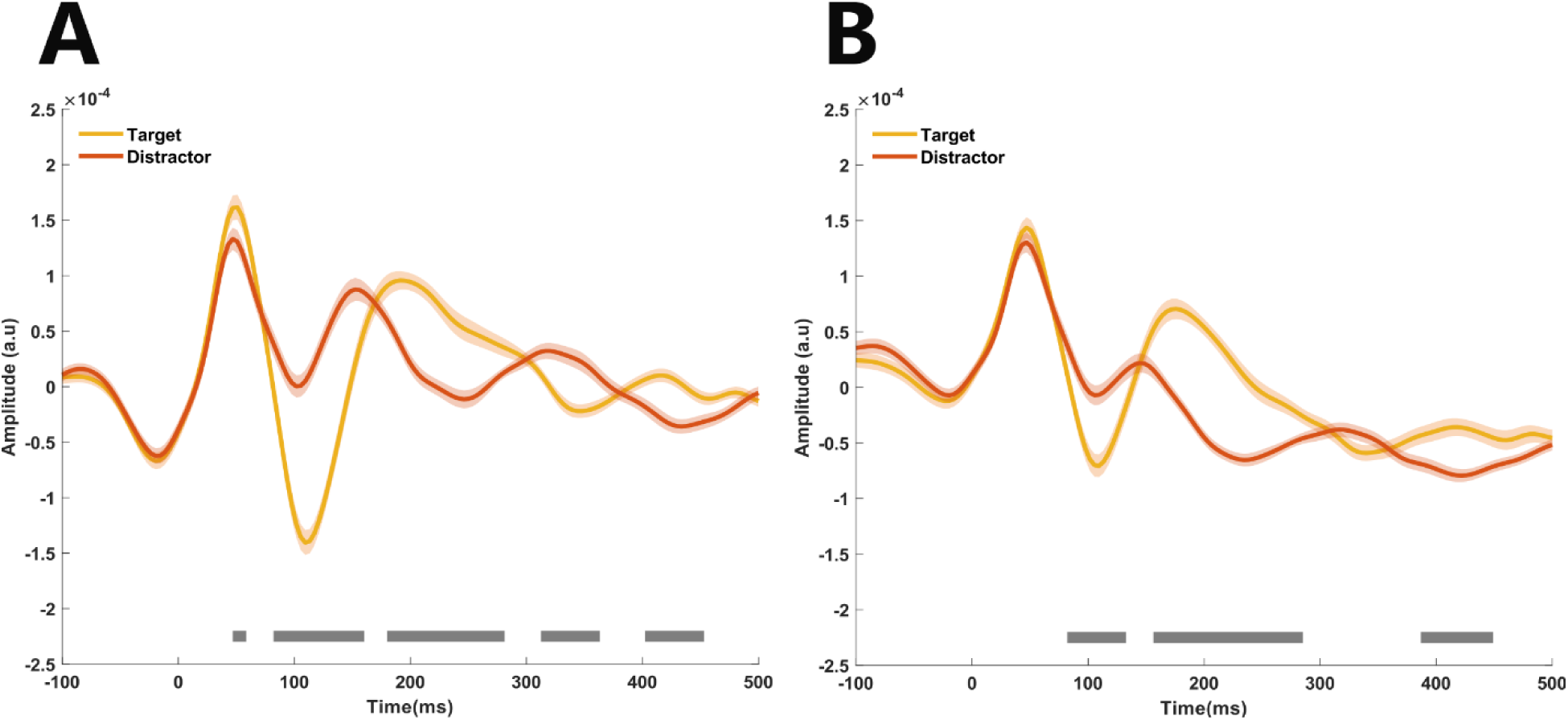
Effect of half-wave rectification of the speech envelope derivative on the TRF. Panel **A** shows the TRFs estimated for target and distracting speech and is identical to Figure 2C. Panel **B** shows the same results, but the speech envelope derivative was half-wave rectified before the TRFs were computed. All other conventions as for Figure 2C.

**Figure S2.**
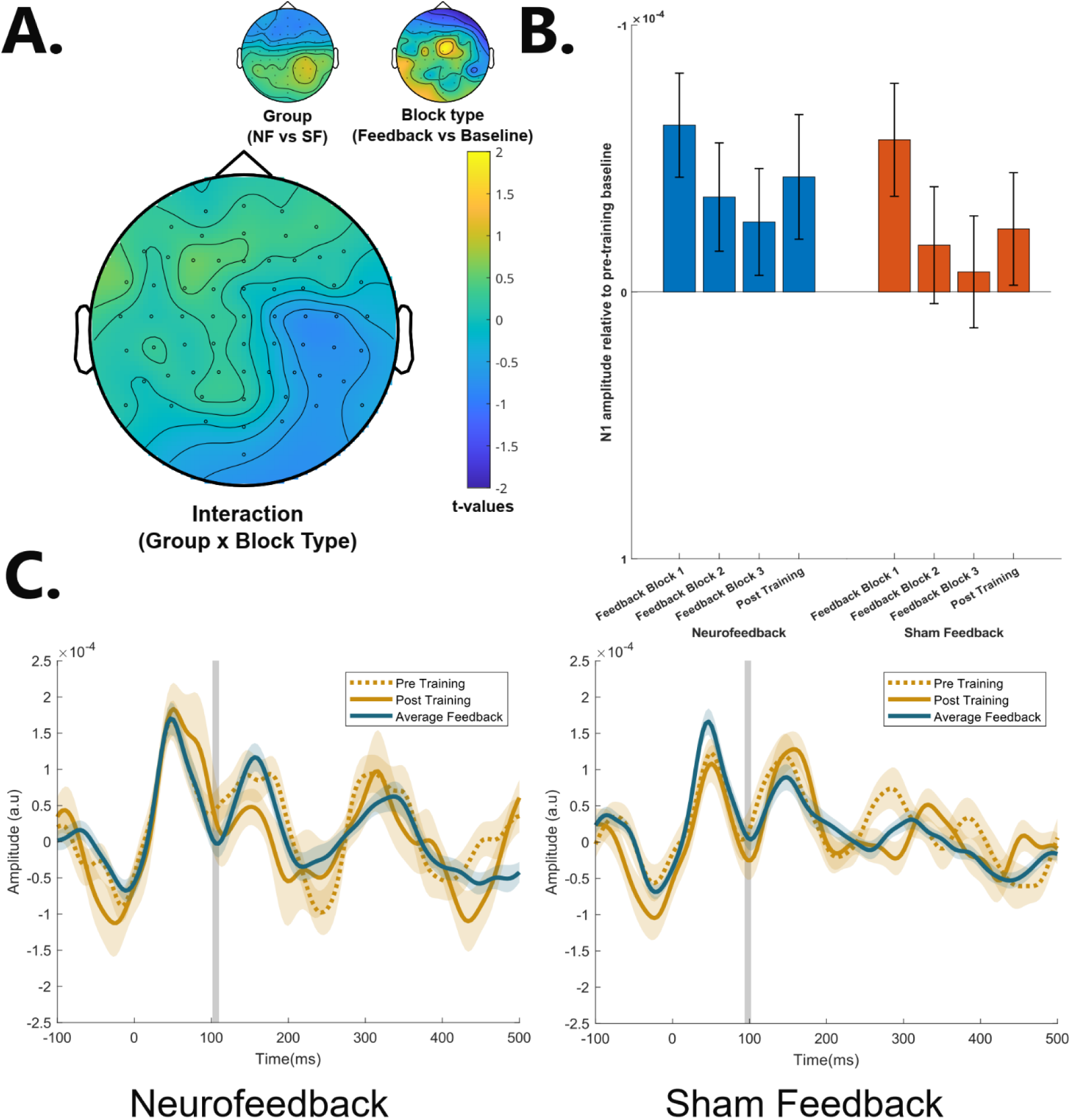
Neurofeedback does not affect the N1 component in the EEG response to distractor speech. **A.** T-values from main effects of group (neurofeedback vs sham feedback) and block type (feedback vs baseline) on N1 amplitudes, as well as their interaction in a linear mixed model. A positive interaction term indicates that the increase in N1 negativity from baseline to feedback blocks is greater in the NF group than in the SF group. B. N1 amplitude values averaged across the nine channels used for feedback, relative to the pre-training baseline (subtraction within participants). Error bars indicate the Standard Error (SEM). C. TRFs estimated from channel FCz, averaged across the three Feedback blocks. Shaded areas around the mean indicate the SEM. Grey shaded area indicates the time window used for statistical analysis.

